# Generation of a human ovarian granulosa cell model from induced pluripotent stem cells

**DOI:** 10.1101/2022.05.15.491993

**Authors:** Dirk Hart, Daniel Rodríguez Gutiérrez, Anna Biason-Lauber

**Affiliations:** Endocrinology Division, Department of Endocrinology, Metabolism and Cardiovascular System, Section of Medicine, University of Fribourg, 1700 Fribourg, Switzerland

**Author notes:** Correspondence (Anna Biason-Lauber).

## Abstract

Sex development is an intricate and crucial process in all vertebrates that ensures the continued propagation of genetic diversity within a species, and ultimately their survival. Perturbations in this process can manifest as variations/differences of sex development (VSD/DSD). Primary gonadal somatic cells – the ovarian granulosa cells (GCs) in the case of women – represent the absolute model to investigate mechanism of disease in VSDs. Collection of these cells in humans is laborious and invasive, while classical animal models fail to recapitulate the human phenotype and function. Furthermore, in patients with the most severe forms of VSD gonadal cells are totally absent. It is therefore vital to develop an alternative cell-model. In view of this, we established an efficient method to reprogram donor-derived urinary progenitor cells (UPs) and differentiate iPSCs into granulosa-like cells (GLCs). The UPs presented a less invasive and high-quality cell source, improving the clinical applicability of the model along with utilising a non-integrative reprogramming method that eliminates alteration of the original genome. This novel GLC model closely resembles human GCs in morphology and marker gene expression of GC cell-fate and essential function. These results provide the prospect to generate patient-specific personalised GC models to investigate mechanism of disease in VSDs and could improve understanding of the intricacies in female gonadal development.

## INTRODUCTION

The healthy development of a female individual rests on the formation of a functional ovary, which in turn depends upon the precursor germ cells that will eventually develop into the female gamete (oocyte) (Eppig, 2018). This transition of primordial germ cells into a mature oocyte requires somatic supporter cells that surround the oocyte in a tightly maintained coexistence between the two cellular compartments of the ovarian follicle. In females, the supporting cell lineage is the granulosa cells (GCs) which can be distinguished into two distinct sub-types: Cumulus granulosa cells (CCs) and mural granulosa cells (MCs). The CCs abut the oocyte membrane and regulate the process of follicle development and directs the differentiation and maturation of the oocyte through dynamic bi-directional cell-to-cell communication (*via* gap junctions) and paracrine signalling events (Binelli and Murphy, 2010). The MCs line the follicular wall creating a layered epithelium with the basal lamina, forming a blood– follicle barrier between the oocyte and theca interna (Gilchrist et al., 2004). Importantly, in the female gonad both populations of GCs drive the development of the endocrine-secreting theca cells that play a vital role during folliculogenesis (Eid and Biason-Lauber, 2016).

Several gonadal disorders are caused by the abnormality of the somatic supporting cells (Hersmus et al., 2008). Therefore, the study of these somatic supporters at a cellular level is crucial for the identification of intrinsic mechanisms central to understanding the pathophysiology of variations/differences of sex development (VSD) and to aid clinical management of infertility. To this goal, a serious impediment is the absence of available patient-specific gonadal cell models.

### A square peg in a round hole

Since the causes of VSD are vastly heterogeneous and complex, an ideal scenario would involve investigation of cells harvested straight from the patient. Through assisted reproductive technologies (ART), such as *in vitro* fertilization (IVF), primary human GCs can be harvested from follicular fluid in the recovered oocyte-GC-complex – but this procedure is invasive for the patient. Furthermore, the long-term *in vitro* cultivation of primary GCs is particularly challenging – as following ovulation GCs become luteinized and are considered terminally differentiated (Niswender et al., 2000). The process of luteinization elicit changes in cell-cell interaction and cell-cycle control – leading to the short proliferative potential observed in culture of mature GCs (Bruckova et al., 2011). Additionally, in some extreme cases of VSD (e.g., patients with Turner syndrome or other primary ovarian failure) these cells are not present at all (Gravholt et al., 2019).

Thus far, animal cell models had been commonly used as alternative for mechanism of disease studies in VSD – but frequently are unable to fully recapitulate the functional effects of detected gene mutations due to species-specific differences in conservation of the sex development network – e.g., in humans, variants of the bone morphogenetic protein 15 (*BMP-15*) gene cause premature ovarian failure (POF), whereas typical follicles form in mice (Di Pasquale et al., 2006; Yan et al., 2001). Over the last decades human granulosa cell lines (such as KGN and COV434) have been established primarily from human ovarian cell tumours (Nishi et al., 2001; van den Berg-Bakker et al., 1993), or immortalised by oncoviruses and oncogenes (Hosokawa et al., 1998; Rainey et al., 1994) – that similarly to animal cell models, have rendered important differences to normal human granulosa cells. Apart from KGN cells, all of these studies could not detect expression of follicle-stimulating hormone-receptor (FSH-R), that is located on GCs in the ovary and is essential for the process of folliculogenesis and estradiol (E2) synthesis in women (Desai et al., 2013). Despite KGN cells being useful to study steroidogenesis and their growth characteristics well-suited to *in vitro* culture, a report found that KGN carried a mutated form of forkhead box L2 (*FOXL2)* 402C>G (C134W) which is linked to progression of human granulosa cell tumours (Cheng et al., 2013). *FOXL2* is regarded as a central regulatory factor in ovarian development and maintenance (Veitia, 2010).

With these considerations in mind, a new source of patient-derived somatic cells that can also be conveniently sampled is fundamental for clarifying patient-specific granulosa cell characteristics and improved clinical applications in gonadal disease.

### An attractive alternative

Cell reprogramming followed by guided *in vitro* differentiation are exciting tools that enable us to harness the pliancy of induced pluripotent stem cells (iPSCs) to generate high-quality *de novo* human stem cell-derived cell sources for utilisation in regenerative medicine and disease modeling (Terryn et al., 2018). Nobel Laureate Shinya Yamanaka and his team first solved the puzzle to convert terminally-differentiated cells via transcriptional reprogramming into cells that are pluripotent (Takahashi and Yamanaka, 2006). These cells share important features of embryonic stem cells (ESCs) and exhibit comparable characteristics of cell-regeneration and the ability to differentiate into a myriad cell-lineages (Buchholz et al., 2009; Burridge et al., 2012). Subsequent studies showed new cell lines that simulate gonadal cells more efficiently compared to previous cell models (Lipskind et al., 2018; Park et al., 2009; Rodriguez Gutierrez et al., 2018).

Herein we set out to generate a human-derived *in vitro* granulosa cell model from iPSCs, using urinary progenitor cells (UPs) as a novel cell source and ectopic expression of reprogramming factors, followed by guided differentiation into granulosa-like cells (GLCs) through a cocktail of growth factors associated with differentiation in GCs. Figure 1 summarizes the stages in the iPSC-derived granulosa cell model generation during this study.

**Figure 1.**
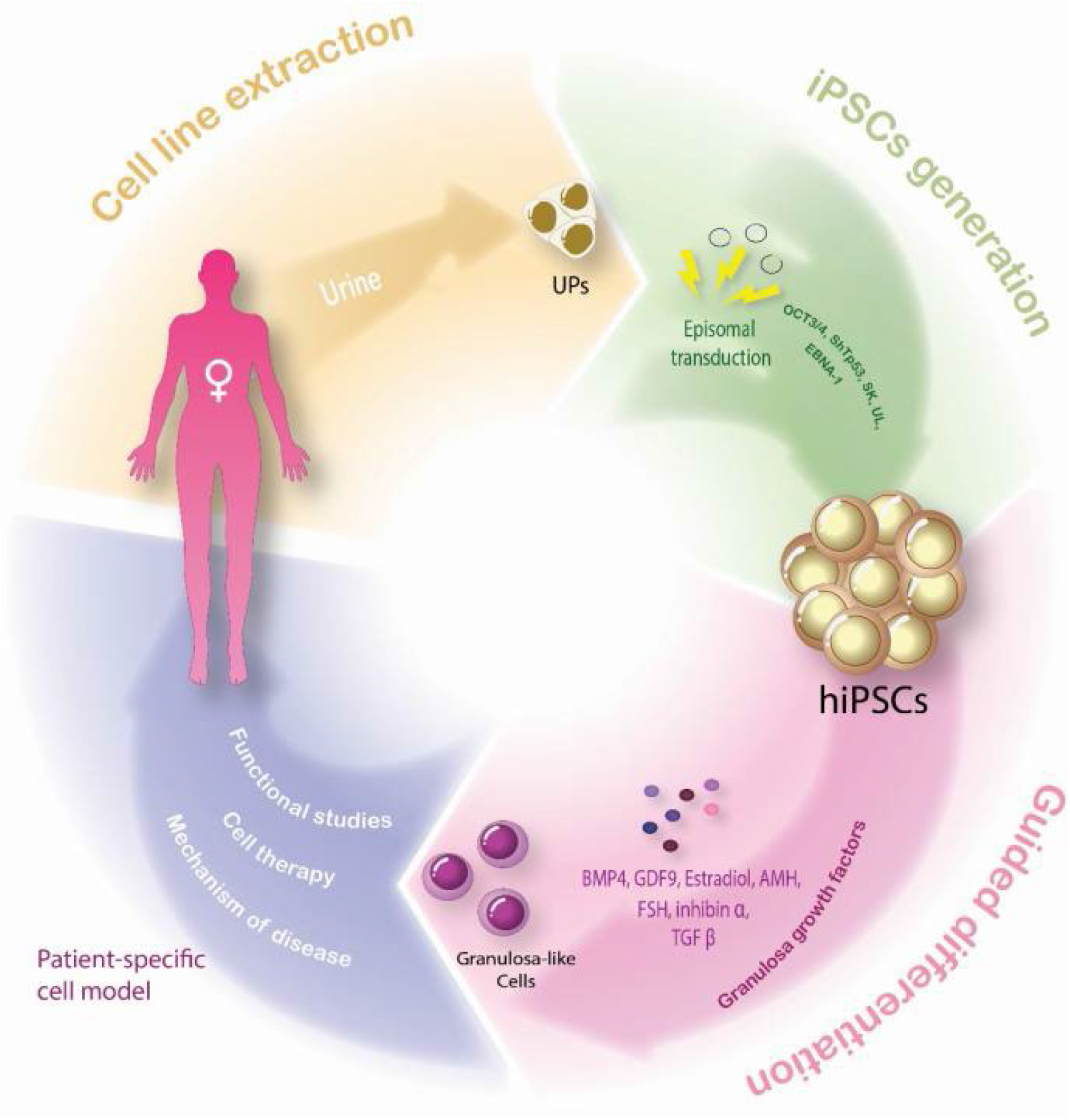
Conceptual schematic representing the four-stage approach to generate a patient-specific cell model of granulosa-like cells by guided differentiation of iPSCs derived from urinary progenitors. FSH, follicle-stimulating hormone; GDF9, growth differentiation factor 9; BMP4, bone morphogenetic protein 4; TGF β, transforming growth factor β. Modified from (Rodriguez Gutierrez and Biason-Lauber, 2019).

## MATERIALS AND METHODS

### Cell line culture

Human ovarian granulosa tumour cells (KGN) (RCB1154, RIKEN BRC) were cultured in Dulbecco’s Modified Eagle’s Medium (DMEM – high glucose; Sigma-Aldrich) and F12 Ham (D6421, Sigma-Aldrich) in a 1:1 ratio, supplemented with 10% Fetal Bovine Serum (FBS; BioConcept Ltd. Amimed) and 1% Penicillin/Streptomycin (Pen/Strep; 15140122 Gibco, ThermoFisher Scientific), and maintained at 37 °C and 5% CO_2_.

### UPs isolation and expansion

Urine was collected from a female human donor, and UP isolation and culture were performed according to previous protocols (Afzal and Strande, 2015; Cheng et al., 2017). Briefly, cells were collected from 100-150 ml of spontaneous urine by centrifugation at 400 x g for 10 min at room temperature (RT). Urine was discarded and the pellet washed with 100 U/ml penicillin, 50 μg/ml streptomycin in 8 ml of phosphate-buffered saline (PBS). After a second centrifugation step at 400 x g for 10 min at ambient temperature, supernatant was discarded and the cells resuspended in UP cell (UPC) medium: an equal mix of Keratinocyte Serum Free medium (KSF, 2101, ScienCell) supplemented with [5 ng/ml epidermal growth factor (EGF), 50 ng/ml bovine pituitary extract (BPE), 30 ng/ml cholera toxin, 1% Pen/Strep] and Progenitor Cell medium consisting of DMEM and F12 Ham (3:1) supplemented with (10% FBS, 0.4 μg/ml hydrocortisone, 0.1 nM cholera toxin, 5 ng/ml insulin, 5 μg/ml transferrin, 0.1 μg/μl triiodo thyronine, 10 ng/ml EGF, 1% Pen/Strep). Cells were then seeded in 24-well plates covered with attachment factor (AF).

### Reprogramming of UPs into iPSCs

Expanded UPs were reprogrammed into iPSCs, based on integration-free nucleofection of episomal reprogramming plasmids, according to manufacturer and prior literature (Afzal and Strande, 2015; Bang et al., 2018) to optimize transfection efficiency and avoid genome-alteration.

UP cells expanded in 6-well culture plates were detached by treatment with 1 ml Trypsin-EDTA (0.25%) per well. Collected cell suspensions were centrifuged 300 x g for 5 min and resuspended in 1 ml D-PBS. Cells were counted with a LUNA-II Automated Cell Counter (Logos Biosystems) and aliquoted to a concentration of 1.75×10^5^ cells/20 μl nucleofection reaction. Using the DC100 electroporation program, nucleofection was performed on the Amaxa 4D-Nucleofector X System (Lonza) in 20 μl Nucleocuvette 16-well strips with the P1 Primary Cell 4D-Nucleofector X Kit, V4XP-1032 (Lonza). A mix of plasmids (all from Addgene) including pCXLE-hOCT3/4-shp53-f (27077), pCXLE-hSK (27078), pCXLE-hUL (27080), and pCXWB-EBNA1 (37624) were added to UP wells at a concentration of 1 μg for each plasmid. Each nucleofection experiment included pmaxGFP Vector (1 μg/μl, Lonza) as positive control for transduction. Electroporated cells were then transferred onto pre-warmed After approximately 12 days expansion, a portion of the cells were harvested for confirmation of UP cell identity by fluorescence activated cell sorting (FACS) analysis. Matrigel-coated wells of a 24-well plate with UPC medium supplemented with 10 µM Y-27632 ROCK inhibitor (STEMCELL Technologies) and incubated at 37°C and 5% CO_2._ Day of nucleofection was considered as Day 0 of the experiment. After 24h (day 1), medium was changed to UPC medium without ROCK inhibitor. At day 3, medium was replaced by 1 ml of a 1:1 mixture of UPC medium and reprogramming medium TeSR-E7 (STEMCELL Technologies). From day 4 to day 18-20, medium was changed every other day.

To identify pluripotency in vivo, generated colonies were stained with Alkaline Phosphatase Live (AP, A14353, Molecular Probes, Life Technologies). AP positive colonies were then manually picked and seeded for further expansion onto Matrigel-coated wells of 24-well plates with mTeSR-1 (STEMCELL Technologies) maintenance medium with 10 µM Y-27632 ROCK inhibitor. Once more, the medium was replaced by mTeSR-1 only, one day after isolation/plating.

### Differentiation of iPSCs into GLCs

GLC differentiation was performed based on insights from previous protocols (Gilchrist et al., 2008; Glister et al., 2003; Jung et al., 2017; Liu et al., 2016), with alterations. Herein, expanded iPSC colonies without spontaneous differentiation were manually selected and seeded in poly 2-hydroxyethyl methacrylate (polyHEMA, Sigma-Aldrich) coated ultralow attachment 6-well plates with 2 ml/well of iPSC medium without growth factors to form embryoid bodies (EBs) during 48-72h. iPSC medium consisted of 1:1 DMEM:F12 Ham supplemented with 10% Knockout serum replacement (KSR, Gibco, ThermoFisher Scientific), 5% non-essential amino acids (NEAA, ThermoFisher Scientific), 1% 2-Mercaptoethanol, 1% Pen/Strep and (10 ng/ml) basic fibroblast growth factor (bFGF, Merck Millipore). Then to induce mesoderm commitment in the EBs, the EBs were transferred to Matrigel-coated wells with iPSC medium supplemented with 50 ng/ml bFGF and 30 ng/ml BMP4 (Gibco, ThermoFisher Scientific) for 24h. After 24h, the iPSC medium [incl. 20 ng/ml EGF (Sigma-Aldrich), and 50 ng/ml bFGF] was supplemented with 20 ng/ml FSH (Sigma) and 20 ng/ml Estradiol (Sigma-Aldrich) for 48h, followed by 20 ng/ml AMH (Sino Biological), 20 ng/ml FSH and 20 ng/ml Estradiol for another 48h. From this point, two different protocols to produce GLCs were used: A) some cells were supplemented with 20 ng/ml AMH, 20 ng/ml Estradiol, 20 ng/ml Inhibin A (ThermoFisher Scientific) and 20 ng/ml Inhibin B (Bio Rad), 15 ng/ml TGF β and 15 ng/ml TGF α (both Gibco, ThermoFisher Scientific) for 96h, or B) the other cells were supplemented with 20 ng/ml AMH, 20 ng/ml Estradiol, 20 ng/ml Inhibin A and B, 25 ng/ml BMP15 (Abbexa) and 50 ng/ml human growth/differentiation factor 9 (GDF9, Cusabio Technology) for 96h. The medium was changed every other day.

From this stage, for long-term cultivation of GLCs, pre-GLCs were cultured using a specific low-FBS medium (Bruckova et al., 2011) consisting of 1:1 DMEM/F12 Ham supplemented with 2% FBS, 1% NEAAs, 0.05 μM Dexamethasone, 100 mM L-glutamine, 0.5% P/S, and supplemented with 20 ng/ml EGF, 20 ng/ml FSH, and 50 ng/ml bFGF.

Figure 2 shows a summary of the stages of UPs cell reprogramming and iPSCs differentiation toward GLCs during this study.

**Figure 2.**
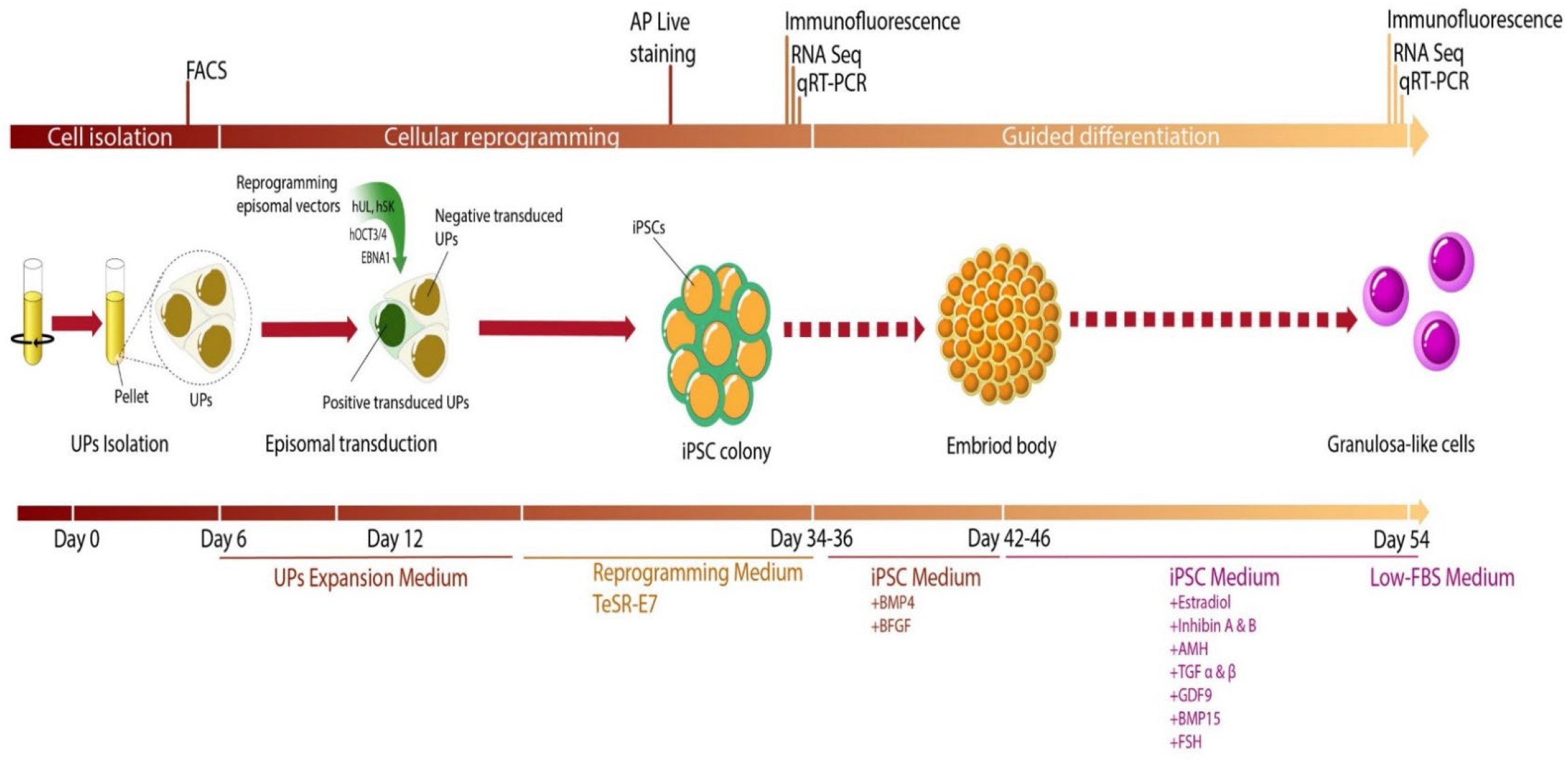
Linear representation of the stages of UPs cell reprogramming followed by iPSCs differentiation toward GLCs.

### Proliferative potential of GLCs

The GLCs were cultured in low-FBS medium as before, and passaged over several days to determine if the cells could be cultivated for long-term. Cells were counted as before, with population doubling time calculated as previously described (Bruckova et al., 2011).

### Immunofluorescence

Pluripotency of iPSC colonies was determined by immunofluorescence (IF). Colonies were fixated with 4% paraformaldehyde (PFA), washed with PBS, and permeabilized by 30 min incubation with 0.2% Triton X-100 in PBS at RT. Colonies were washed with PBS, then blocked with 3% BSA in PBS for 1h, washed 3 times in PBS, and then incubated with primary antibodies against OCT3/4 (1:200, BD Bioscience, 611203), SOX2 (1:500, Abcam, ab97959), TRA-1-81 (1:200, Abcam, ab16289), SSEA1 (1:100, DSHB, MC-480), SSEA3 (1:3, DSHB, MC-631), and SSEA4 (1:3, DSHB, MC-813-70) overnight at 4°C. For Granulosa-like assessment, primary antibodies against FOXL2 (1:500, LifeSpan Bioscience, LS-B12865), and FSH-R (1:100, Abcam, ab113421) were incubated overnight at 4°C. Secondary antibodies Alexa Fluor 488 conjugated goat anti-rabbit (A-11008), Alexa Fluor 594 conjugated goat anti-mouse (A-11005), and Alexa Fluor 488 conjugated donkey anti-goat (A-11055) (all from ThermoFisher Scientific) were diluted 1:1000 and incubated for 1h in the dark at RT. DAPI (Vectashield) mounting medium was added and incubated for 20 min at RT, then samples were inspected under IF on an ECLIPSE Ni-U upright microscope (Nikon).

### AMH quantification

KGN, UPs and GLCs cultured as before in 12-well plates, were washed with PBS and then incubated with Opti-MEM Reduced Serum Medium for 2h.

### Karyotype analysis

Giemsa-banded karyotyping was performed for the UP and GLC clones. Cells were treated with 0.1 μg/ml KaryoMAX Colcemid Solution (15210040, ThermoFisher Scientific) and incubated for 30 min at 37°C, washed with PBS, then dissociated by Trypsin-EDTA (0.25%) and incubated with pre-warmed hypotonic solution (0.075 M KCl) at 37°C for 15 min. Cells were then fixed in methyl alcohol/glacial acetic acid 3:1 and kept at 4°C. At least 10 metaphases were karyotyped by the cytogenic laboratory at the University Children’s Hospital Zurich (Switzerland).

Cell supernatants were collected by trypsinization and cells counted as previously. Secreted AMH in cell culture supernatants was assessed using a microplate Enzyme-linked Immunosorbent Assay (ELISA) kit (ABIN6574075, Antibodies-Online GmbH) and performed according to the manufacturer’s protocol. Results were analysed using an Infinite microplate reader (Tecan Trading AG). Values were normalised according to the number of cells present in each sample.

### Estradiol quantification by LC-MS/MS

Upon preparation of GLC, UP and KGN samples for Estradiol (E2) quantification, measurements were performed in collaboration by the Christoph Seger laboratory at Dr. Risch (Buchs SG, Switzerland). Estradiol was quantified in an ISO 17025 accredited laboratory environment with a liquid chromatography tandem mass spectrometry (LC-MS/MS) assay dedicated for the analysis of steroid hormones in human serum. Matrix exchange experiments did prove the assay was suitable for quantification in cell culture matrix. Prior to analysis, samples (100 µl culture broth) were subjected to protein precipitation followed by phospholipid removal (HybridSPE, Merck/Sigma Aldrich). The internal standard E2-d_5_ (Chromsystems) was added before the precipitation step. A 40 µl aliquot of the purified sample was used for analysis with the LC-MS/MS system (Sciex API6500+ triple quad mass spectrometer hyphenated to an Agilent 1290 Infinity II UHPLC). Chromatographic separation was performed on a biphenyl stationary phase (Restek Raptor Biphenyl 100×2.1, 2.7 µm, BGB Analytik). Gradient elution (water/methanol with 0.2 mM NH_4_F as additive) resulted in E2/E2-d_5_ retention time of 2.6 min. Analyte detection was performed in the negative ESI mode with mass transitions m/z 271→145 for E2, 276→145 for E2-d_5_ for quantification, and 271→183 as confirmative E2 transition. A linear 1/x^2^ weighted calibration function (range 15-18900 pM, eight levels) based on an IVD-CE certified and traceable calibrator system (Chromsystems) was utilized for E2 quantification. Measuring IVD-CE certified quality control materials (Chromsystems) and certified reference materials (BCR-576, BCR-577, BCR-578, IRMM, Merck/Sigma Aldrich) over five days, the E2 assay showed an intermediate precision of 6% (RSD) and an accuracy of ± 7%.

### RT-qPCR

A battery of pluripotency markers and classical granulosa cell markers (see supplemental Table S1 for primers) were used to analyse their expression in the GLCs relative to the values of KGN and UPs. Total RNA was extracted from UPs, KGN, and GLCs using the RNeasy Plus Mini Kit (74136, Qiagen) and 2 μg of extracted total RNA was reverse-transcribed using the Omniscript RT Kit (205111, Qiagen) according to manufacturer’s instructions, respectively. All samples were run in triplicates, with a -RT and no template control (NTC) included in every run. Experiments were performed on a StepOnePlus Real-Time PCR system (ABI, Life Technologies) and PCR products were quantified fluorometrically using the KAPA SYBR FAST qPCR Master Mix (2X) Kit (KR0389, KAPA Biosystems).

To determine the expression level of selected targets, the comparative threshold cycle (Ct) method was applied, using the reference mRNA gene cyclophilin (PPIA) to normalize the data.

### Single-cell RNA sequencing and bioinformatic analysis

Single-cell RNA sequencing (scRNA-seq) was performed at the Functional Genomics Center Zürich (FGCZ) by Chromium 10x sequencing of four samples (UPs, KGN cells, GLC clone A, and GLC clone B) on a Chromium Controller instrument (10xGenomics). 10,000 cells were sequenced for each sample, with the quality metrics of raw reads controlled using FastQC (Babraham Bioinformatics). Alignment of reads and barcode counting were processed on Cell Ranger (10xGenomics), and contamination screening of reads was done with FastQ Screen (Babraham Bioinformatics). For cell filtering and aggregated cell-type clustering analysis, data of the samples was concatenated and normalized, resulting in a combined gene-barcode matrix of all samples. Further filtering and clustering analyses were performed with the Scanpy (v.1.8.2) Python package as previously described (Wolf et al., 2018). For each aggregated data set, cells were normalized into counts per million reads (CPM) and log-transformed (Log (CPM+1)). Low quality cells (fragmented or doublets) were excluded from the analysis by filtering out cells with greater than 7,000 and fewer than 1,000 genes expressed, as well as all those with a percentage of mitochondrial genes greater than 5%. Highly variable genes in the data set (considering those genes with a mean expression range of 0.0125 - 6.0 and minimum normalized dispersion of 0.5) were used as input for principal component analysis (PCA). Uniform manifold approximation and projection (UMAP) dimensionality reduction and shared nearest neighbour clustering (SNN) were applied for clustering and representing populations in a two-dimensional space.

### Statistical analysis

Unpaired t-test analysis was performed using GraphPad Prism 8.4.3 (GraphPad), and all data are given as mean ± SD (Standard Deviation) from at least 3 biological replicates. P<0.05 was considered to indicate a statistically significant difference.

## RESULTS

### UP culture and characterization

Urine was collected from one female human donor, and UP isolation and culture were performed according to previous protocols (Afzal and Strande, 2015; Cheng et al., 2017). Other cell populations present in the urine (such as squamous cells and the occasional blood cell) did not attach and were removed with media changes. After 4 days, small colonies of 3-5 cells were observed and UPs expanded rapidly, reaching confluency in 12 days (Figure 3 A-C). The major proportion of the cells were CD44^+^/CD73^+^/CD146^+^ which are typical markers consistent with their uroepithelial progenitor origin (Figure 3 D) (Bharadwaj et al., 2011; Lang et al., 2013).

**Figure 3.**
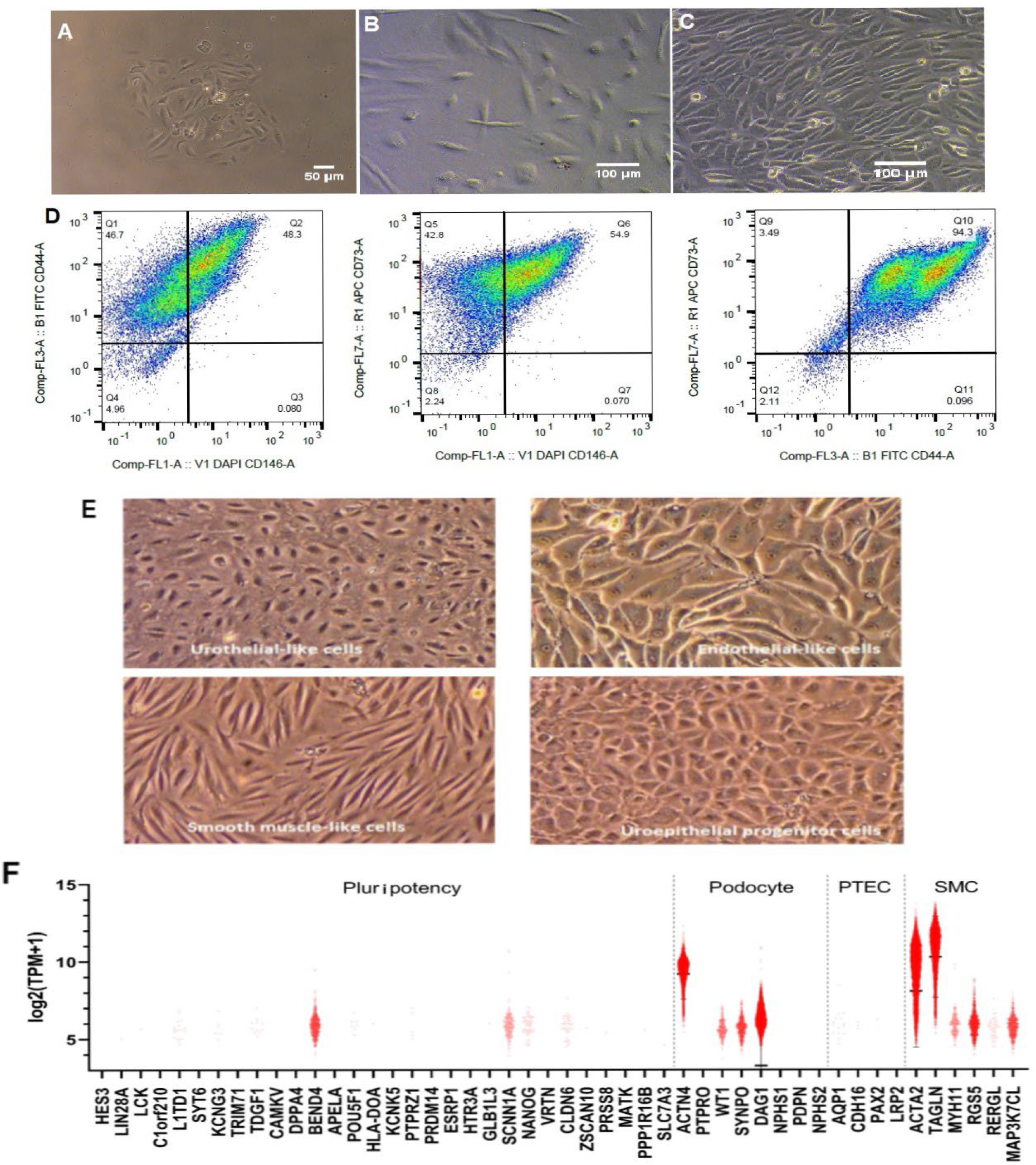
Characterization of UPs and spontaneous differentiation of UPs. UPs were isolated from urine by centrifugation and cultured in UPC medium. **(A)** Cell morphology of UPs on day 4 after isolation. **(C)** Day 12 after isolation. **(D)** FACS analysis showed that 94.3% of cells were CD73+/CD44+ positive, 54.9% expressed CD73+/CD146+, and 48.3% CD44+/CD146+, which confirm their origin as uroepithelial progenitors. **(E)** Phase contrast microscopy of urine derived cells in culture showing multiple cell lineages: Urothelial-like, endothelial-like, smooth muscle-like, uroepithelial progenitor cells. **(F)** Violin plots generated from scRNA-seq data analysis indicating the relative expression levels of marker genes for different cell-types constituting a UP cell population: Pluripotency genes, podocytes, renal proximal tubular epithelial cells (PTEC), and smooth muscle cells (SMC). Plots represent gene expression within the total UP cell population analysed. Expression level represented as normalized log2 Transcripts Per Million +1 (log2 [TPM+1]).

### UPs have potential to spontaneously differentiate

Urine derived cells have been shown to possess the capacity for differentiating into a myriad of cell lineages (Bharadwaj et al., 2013; Lazzeri et al., 2015; Liu et al., 2018; Wan et al., 2018). During culture of UPs in our setting a portion of cells within the UP population spontaneously differentiated into Urothelial-like, endothelial-like, smooth muscle-like, and uroepithelial progenitor cells (Figure 3 E). Marker genes expressed for cell-types typical of differentiated UP cell populations were analysed from our scRNA-seq data, confirming differentiation of some UP cells from a pluripotent state into a mix of renal cell-types (Figure 3 F).

### UPs are reprogrammed into iPSCs

UPs were reprogrammed into iPSCs, according to manufacturer and prior literature (Afzal and Strande, 2015; Bang et al., 2018). Approximately 18 days post nucleofection, colonies typical in morphology of iPSCs appeared: small cells with defined borders and high nuclear/cytoplasmic ratio (Kato et al., 2016) (Figure 4 A). Pluripotency of iPSC colonies in vivo was determined by alkaline phosphatase live staining (Figure 4 A) and IF of pluripotent cell markers was assessed (Figure 4 B). The pluripotent cell markers OCT3/4, SOX2, SSEA3, SSEA4 and TRA-1-81 were clearly expressed in the nuclei of colonies compared to SSEA1, a marker of differentiation which could not be detected (Figure 4 B). This indicates the UPs were efficiently reprogrammed into iPSCs.

**Figure 4.**
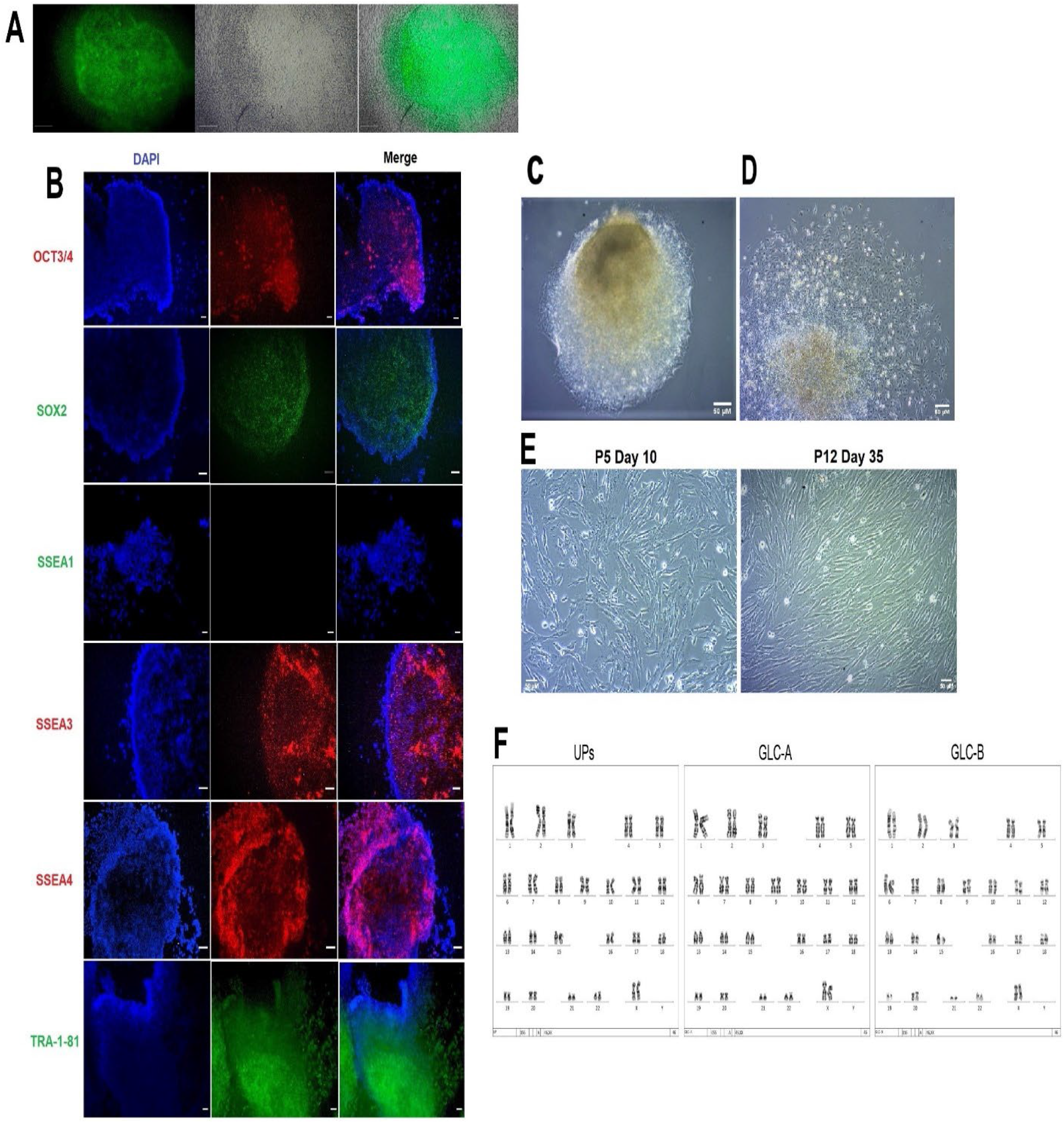
iPSCs characterization by IF and guided differentiation of iPSCs to Granulosa-like cells. **(A)** Pluripotency was assessed by alkaline phosphatase live staining. **(B)** Colonies were stained for antibodies against the pluripotent markers OCT3/4, SOX2, SSEA3, SSEA4 and TRA-1-81. Mounting medium contains DAPI stain. SSEA1 differentiation marker was used as negative control. **(C)** Embryoid body growing in suspension. **(D)** Embryoid body in differentiation, the undifferentiated center (light brown) can be observed while cells on the periphery have changed in morphology. **(E)** Long-term cultivation of established GLCs with characteristic spindle-shaped (fibroblast-like) morphology of human Granulosa cells. **(F)** Representative G-banding karyotype images of control UP cells and GLC clones (A and B). Analysis of 10 metaphase cells for each sample demonstrated a normal female karyotype (46,XX) in all samples, with no chromosomal aberrations reported. P, passage. Scale bars 50 μM.

### Guided differentiation of iPSCs into GLCs

Granulosa-like cell differentiation was performed based on insights from previous protocols and guided by a modified cocktail of extracellular signalling molecules with known function in GC formation, including AMH, Estradiol, TGF α & β, BMP15, and GDF9 (Gilchrist et al., 2008; Glister et al., 2003; Jung et al., 2017; Liu et al., 2016). A critical step in embryonic stem cell differentiation is the formation of embryoid bodies (EBs) (Doetschman et al., 1985). The iPSC colonies grew in suspension to effectively form EBs and were induced to mesoderm commitment by addition of BMP4 and bFGF to the culture medium (Figure 4 C). During guided differentiation, cells on the border of the EBs transitioned into the typical spindle-shaped morphology (fibroblast-like) of human GCs (Ai et al., 2019) (Figure 4 D). After approximately 12 days the GLCs reached confluency and were able to be subcultured over several passages for longer than 30 days using a specific low-FBS medium (Bruckova et al., 2011) (Figure 4 E). The proliferative activity of the GLCs were comparable to the KGN cell line, with the average doubling time of the GLCs being 21.9 ± 5.7 h and 22.8 ± 10.7 h for KGN (data not shown).

### GLCs present an unaltered genome

As quality control screening Giemsa-banded karyotyping was performed for the UPs and GLC clones to ensure the genome of parental cells is maintained in an uncompromised manner. Karyotype analysis revealed there were no chromosomal abnormalities in the GLC clones compared to UP cells (Figure 4 F), indicating good genomic integrity of the GLCs.

### GLCs express the Granulosa cell markers FOXL2 and FSH-R

The GC markers FOXL2 (master regulator gene) and FSH-R play critical roles in regulating the ovarian reserve and follicular development (Desai et al., 2013; Uhlenhaut and Treier, 2011), and were assessed in the GLC clones, UPs (negative control line) and KGN (positive control line) by immunofluorescence. FOXL2 was clearly detectable in all GLC clones and KGN cells, and not present in UPs (Figure 5 A). FSH-R was present in all GLC clones and seemed more highly expressed compared to KGN, and not present in UPs (Figure 5 B).

**Figure 5.**
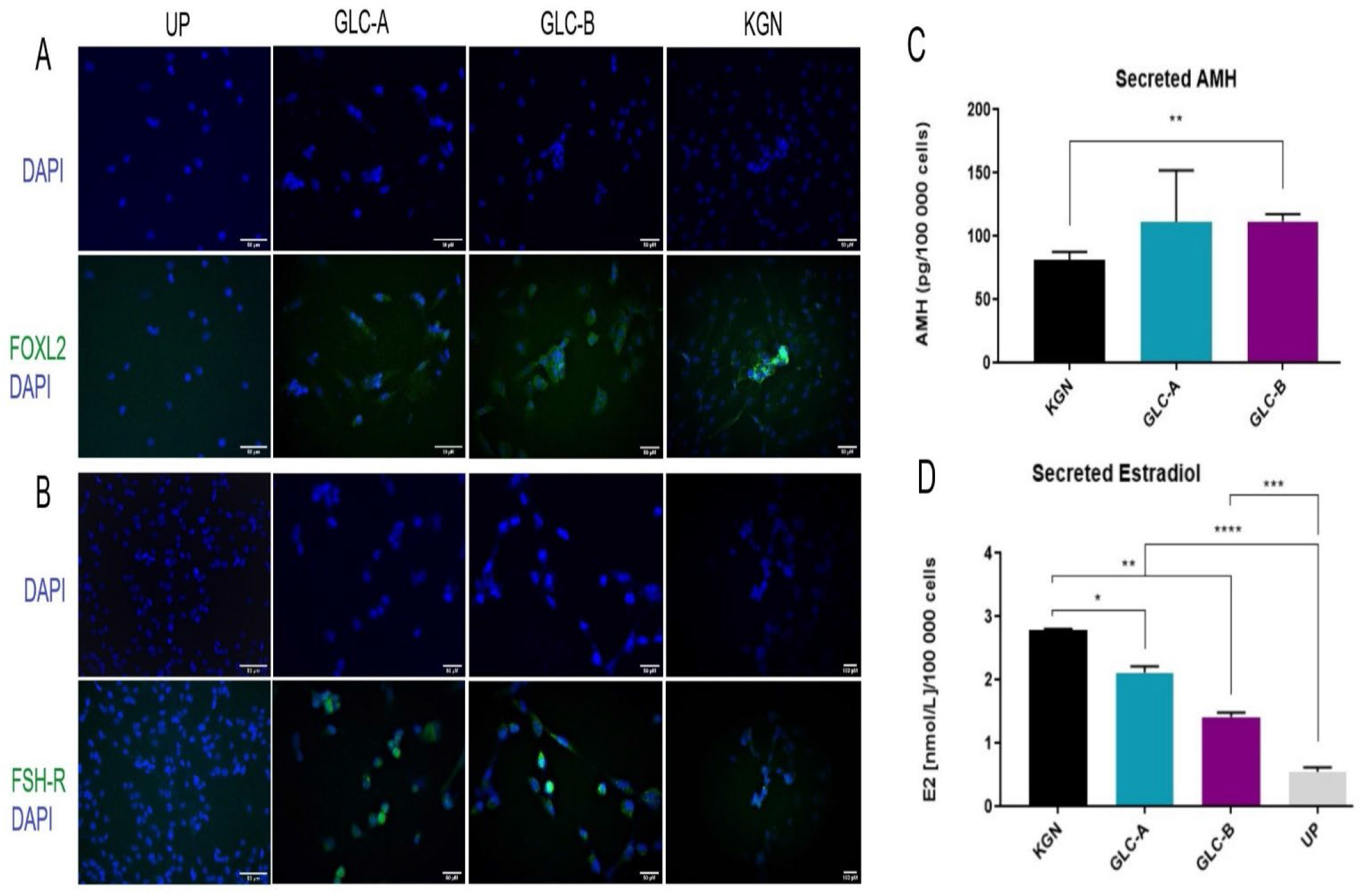
Immunofluorescence staining of GLC clones and functional characterisation of hormone and steroid production in the GLCs. **(A)** UPs, GLC clones (A and B) and KGN cells stained for antibodies against FOXL2. **(B)** UPs, GLC clones (A and B) and KGN cells stained for antibodies against FSH-R. Mounting medium contains DAPI stain. Scale bars 50 μM. **(C)** Secreted AMH quantification by ELISA. Note: Since the ELISA could not detect any levels of AMH in UPs, no statistical values for UPs could be included on the graph. **(D)** Secreted Estradiol measured by LC-MS/MS. The relative values were compared to UPs and KGN cells and normalised per 100 000 cells. Three replicates were conducted for all experiments and significant change determined utilizing the unpaired t-test (n =3). All data are given as mean ± SD (error bars). For the significance: ****, P < 0.0001; ***, P < 0.001; **, P < 0.01; *, P < 0.05; non-significant differences are not indicated. AMH, Anti-Müllerian hormone; pg, picogram; E2, Estradiol; nmol/L, nanomole per liter.

### GLCs secrete AMH and produce Estradiol

Anti-Müllerian hormone (AMH) is a well-known marker of granulosa cells (Depmann et al., 2015). AMH secreted in the medium of GLCs was measured by ELISA and compared with AMH levels in UPs and KGN cells. Relative to the KGN cells, the levels of secreted AMH were higher in both GLC clones (Figure 5 C). GLC-A was not significantly higher (P = 0.2714), but GLC-B had significantly elevated levels of AMH (P = 0.0035). AMH was not detected in the medium of the UPs.

The major steroid hormone produced by the preovulatory GCs in women is estradiol (E2) (Ryan, 1982). E2 (17α-Estradiol) secreted in the medium of GLCs was quantified by LC-MS/MS relative to E2 levels of the known steroidogenic cell line KGN and UPs (see Supplemental Figure S1 for representative measurements). Both GLC clones were able to produce high amounts of E2 (i.e., >= 0.073-1.101 nmol/L for premenopausal women (Stanczyk and Clarke, 2014)). Compared to KGN, E2 secretion levels in the GLC clones (GLC-A and GLC-B) were lower, but significantly higher than the marginal levels measured in UPs (Figure 5 D).

### Global patterns of single-cell expression profiles

UMAP dimensionality reduction and SNN clustering could represent the four cell populations analysed in a two-dimensional space, with both GLC clones (GLC-A and GLC-B) mostly overlapping and clear spatial separation between UPs, KGN, and GLCs (Figure 6 A). Transcriptome analysis based on total gene expression confirmed the highly similar expression profiles of the GLC clones with no significant differences observed (Figure 6 B), suggesting that the differentiation protocol is reproducible. GLCs had a partly-shared expression pattern with KGN, but a diverse expression profile compared to UPs (Figure 6 B), which indicates the GLCs were efficiently differentiated from UPs.

**Figure 6.**
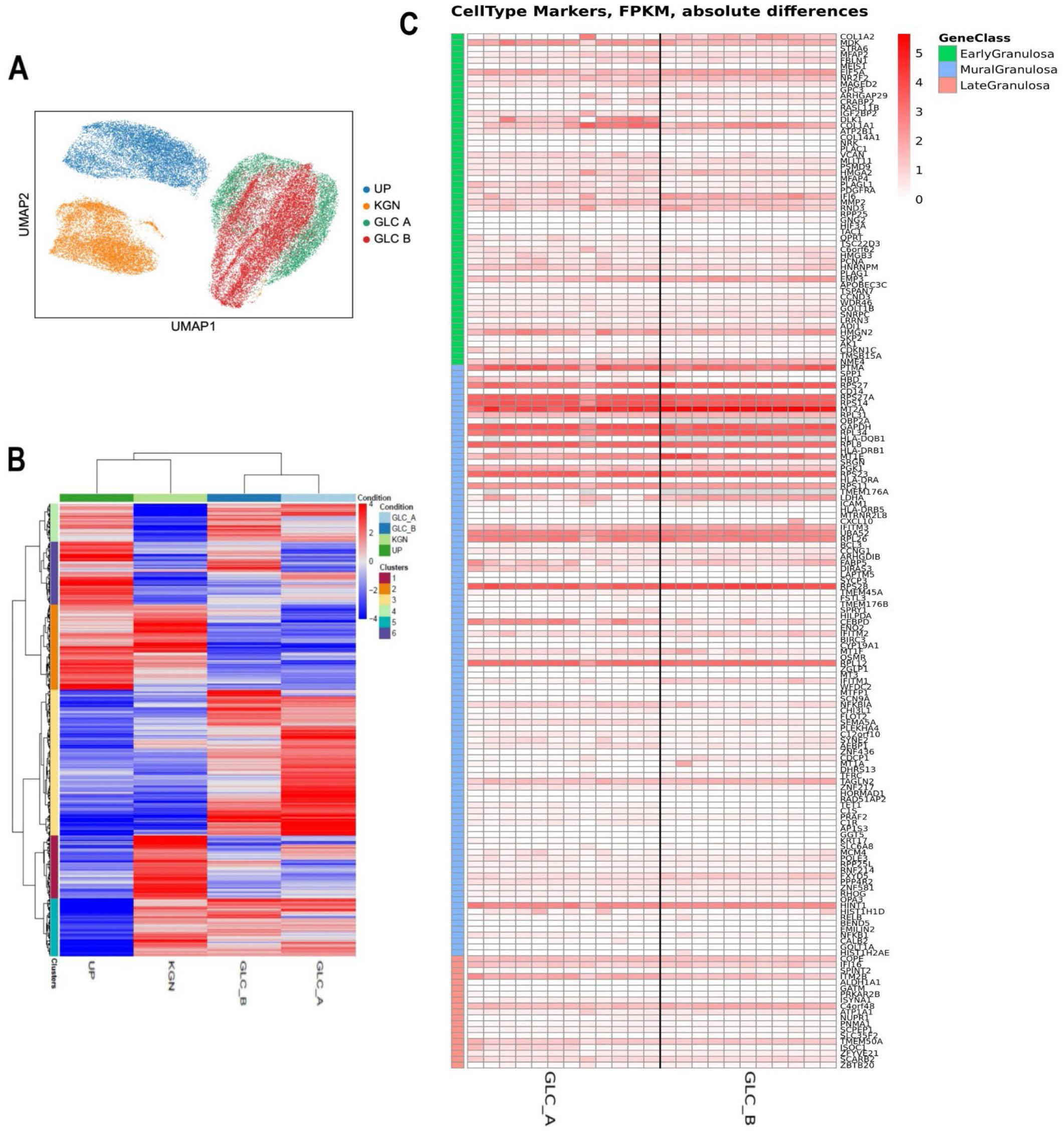
Classification of cell populations by total gene expression and cell state of the GLC clones based on granulosa stage-specific cell marker expression. **(A)** Cell-type cluster separation by UMAP dimensionality reduction. Cells colored by the identified cell-types. **(B)** Transcriptome heatmap of total genes expressed in the four cell populations analysed (UPs, KGN, GLC-A, GLC-B). Red color means higher expression and blue color means lower expression than the average. **(C)** Heatmap of GC marker genes grouped according to GC cell-stage (early, mural, late granulosa) and hierarchical expression. Red color means higher expression, white color means slight or no expression. Expression pattern is of grouped cell clusters for GLC-A and GLC-B. Expression level represented as Fragments per Kilobase Million (FPKM).

### Global patterns of single-cell expression profiles

UMAP dimensionality reduction and SNN clustering could represent the four cell populations analysed in a two-dimensional space, with both GLC clones (GLC-A and GLC-B) mostly overlapping and clear spatial separation between UPs, KGN, and GLCs (Figure 6 A). Transcriptome analysis based on total gene expression confirmed the highly similar expression profiles of the GLC clones with no significant differences observed (Figure 6 B), suggesting that the differentiation protocol is reproducible. GLCs had a partly-shared expression pattern with KGN, but a diverse expression profile compared to UPs (Figure 6 B), which indicates the GLCs were efficiently differentiated from UPs.

### Dynamic gene expression profile of GLCs

Granulosa cell population type varies according to the stages of folliculogenesis (Gougeon, 2010). In order to determine the cell state of the GLCs, we assessed our GLC scRNA-seq data against a reference set of granulosa stage-specific (i.e., early, mural, late granulosa) marker genes extracted from previous scRNA-seq data on human gonadal niche cells (Li et al., 2017). The GLC clones (GLC-A and GLC-B) seemed to have a gene expression profile predominantly corresponding to mural GCs (in number of marker genes expressed and level of expression), but also expressed several marker genes of early-stage GCs and to a lesser extent some markers of late GCs (Figure 6 C). This suggests a dynamic granulosa cell state within the overall cell population of the GLCs, which most likely reflects the varying degree of differentiation in individual granulosa-like cells.

### GLCs express classical markers of human granulosa cell fate and function

To further ascertain if the GLCs are indeed granulosa-like, we selected a set of classical marker genes shown to be involved in Granulosa cell fate (GC proliferation and differentiation) and function (folliculogenesis and steroidogenesis) (Feuerstein et al., 2007; Georges et al., 2014; Grondahl et al., 2012; Hamel et al., 2008; Li et al., 2017; Miller and Auchus, 2011; Noma et al., 2011; Stocco, 2008; Zhang et al., 2018). The total expression of GC marker genes was compared for UPs, KGN, and the GLC clones within our scRNA-seq data set (Figure 7 A), and was validated by corresponding analysis of gene expression relative to iPSCs by RT-qPCR (Figure 7 B). The expression levels of all pluripotency markers analysed (NANOG, OCT4, SOX2, DPPA2, and DPPA4) were lower in all cell populations relative to the iPSCs, especially so in the GLCs (Figure 7 B), which additionally confirms successful differentiation away from iPSCs. Overall, the GLCs showed expression of most GC-specific markers analysed, specifically important markers of cell proliferation and differentiation (FOXL2, INHBA, AMH, TST, KITLG, FST, CDCA3, NRG1), steroidogenesis (CYP19A1, STAR, AR, HSD17B1, HSD3B2, ESR2, PGR, FSH-R, LH-R), and granulosa dependent folliculogenesis (FOXL2, AMH, ZEB2, CD44, KDSR, HSPG2, CHST8) (Figure 7 A and B). When compared to the KGN control cell population, the GLCs had in general similar expression levels of granulosa marker genes, although the expression level of some markers in the GLCs were highly elevated (HSD17B1, HSD3B2, FST, FSH-R). Interestingly, we observed UPs also expressed several of the genes analysed, and noted FOXL2 expression levels in KGN and UPs were slightly higher than the GLCs.

**Figure 7.**
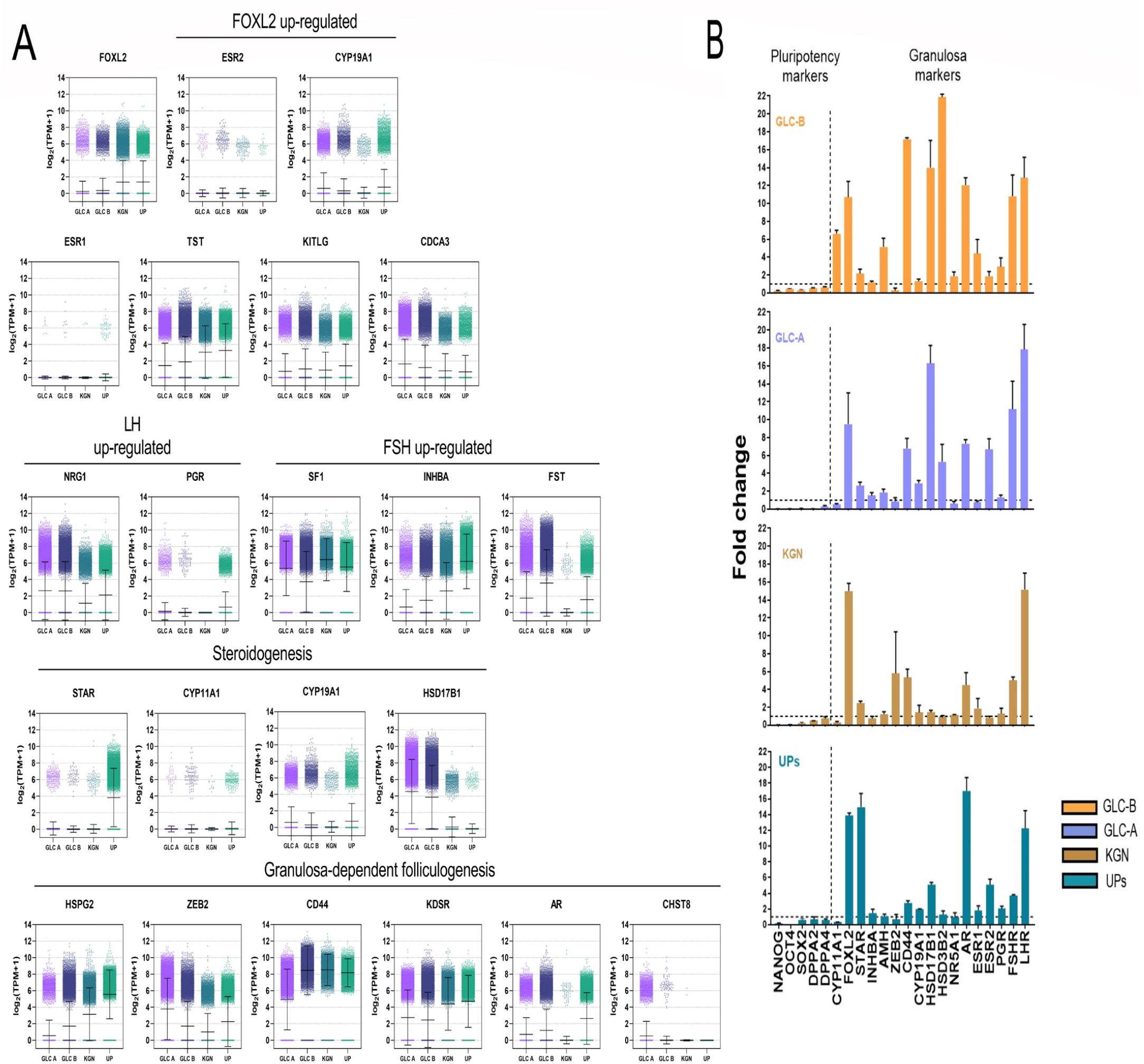
Gene expression analysis of granulosa cell-specific markers. **(A)** Scatter dot plots showing the total expression level of relevant marker genes reflecting human granulosa cell-specific function and fate. Plots represent gene expression within the total cell population of UPs, KGN, GLC-A and GLC-B. Expression level represented as normalized log2 Transcripts Per Million +1 (log2 [TPM+1]). All data are given as mean (middle bar) ± SD (error bars). **(B)** RT-qPCR data of classical markers involved in human Granulosa cell differentiation and function. Comparative analysis between UPs, KGN, GLC-A and GLC-B. All values are relative to iPSCs expression level (dotted horizontal line (fold change =1)). For expression experiments, three replicates were conducted and significant change determined utilizing the unpaired t-test (n =3). All data are given as mean ± SD (error bars). LH, luteinizing hormone; FSH, follicle stimulating hormone; FOXL2, forkhead box L2.

## DISCUSSION

In the developing female gonad granulosa cells function as the main supporting niche for precursor germ cells and facilitates the development of these germ cells into a mature oocyte – which is ultimately essential for the formation of a functional ovary and the ability to reproduce. Numerous developmental divergences such as VSDs arise from anomalies in GCs. A major obstacle to an improved understanding of the genetic programs and molecular mechanisms that lead to disease in VSDs is the lack of a suitable model system for the single patient. The invasive nature of collecting human luteinized GCs (that are terminally differentiated) from IVF procedures – coupled with their problematic cultivation and short lifespan *in vitro* – and moreover the fact patients with VSD due to gonadal dysgenesis do not have any gonadal cells at all necessitated the search for different cell sources to generate a sufficient model.

Previous cell models acquired from animals are often not fully appropriate for studying the functional consequences of discovered variants, and models derived from tumorigenic ovarian cells (e.g., COV434 and KGN) are characterised by key differences to healthy human GCs and mutations in important female-specific developmental factors (FOXL2 in KGN). Technological advances, such as the ability to induce a pluripotent state in terminally differentiated cells through cellular reprogramming and thereafter guided *in vitro* differentiation into multiple self-renewing cell types, precipitated exciting novel cell sources for applications in regenerative medicine and drug discovery. This prompted us to take advantage of these versatile technologies to generate a human donor-derived granulosa cell model from iPSCs, which could be utilized amongst others to improve the understanding of the mechanisms that lead to disease in VSD patients.

Common cell sources for reprogramming have been derived from somatic cells like fibroblasts or peripheral blood mononuclear cells (PBMCs) (Simara et al., 2018; Takahashi and Yamanaka, 2006). Fibroblasts are terminally differentiated and uncomplicated to culture *in vitro*, however skin biopsy is required and repeated subcultures to have sufficient cells for reprogramming (Yu et al., 2007). PBMCs can be acquired from routine venous blood sampling, but reprogramming of non-adherent cells is notoriously challenging with poor efficiency less than 0.01% reported (Kim et al., 2016; Loh et al., 2009). Uroepithelial progenitor cells have been shown to differentiate with high efficiency into the three dermal cell lineages expressing smooth muscle, endothelial and urothelial cell markers, while also having telomerase activity and being immensely expandable but without the formation of tumours or teratomas *in vivo* (Liu et al., 2018). UPs are more swiftly reprogrammed into iPSCs and with improved efficacy compared to fibroblasts or other mesenchymal cells (Guan et al., 2014).Therefore, as alternative cell source in this study we isolated UPs from urine using a non-invasive, donor-derived, easy, and low-cost approach. To our knowledge, this is the first report on the use of UPs for the purpose of generating granulosa-like cells.

A further concern in reprogramming experiments is the generation of integration-free iPSCs. Systems based on synthesised mRNA technology or using Sendai virus vectors have been employed towards this purpose (Fusaki et al., 2009; Warren et al., 2010), but alteration of mRNA is a complex process and altered-mRNA lasts a very short duration in host cells, while Sendai virus experiments are costly and require strict biological safety measures (Fusaki et al., 2009; Mandal and Rossi, 2013). In our system, we were able to rapidly and efficiently reprogram expanded UPs into iPSCs exhibiting excellent pluripotency, and moreover by utilising an integration-free methodology based on nucleofection of episomal vectors (Afzal and Strande, 2015; Bang et al., 2018) in order to optimize transfection efficiency and maintain the original genome of human-derived cells (UPs) – as evidenced by no chromosomal alterations reported in the karyotype analysis of the GLCs generated in this study from iPSCs derived through reprogramming UPs.

Guided differentiation of iPSC colonies into different cell types can be achieved by supplementing growth medium with specific extracellular signalling factors in a time-exact manner – the selection of which are essential for differentiation into the targeted cell-type (Gunhanlar et al., 2018; Pandya et al., 2017). Our protocol used growth factors associated with female gonadal development and GC differentiation: BMP4 and bFGF induces mesoderm commitment which is critical during the development of the urogenital system, while EGF and bFGF is known to aid proliferation of primary GCs in culture (Bruckova et al., 2011; Evseenko et al., 2010; Lawson et al., 1999). Inhibin-alpha and inhibin-beta which are secreted by GCs play a vital role in the maintenance of ovarian follicular development and maturation (Bao et al., 2021; Myers et al., 2009). The final step in our differentiation protocol involved addition to the medium of oocyte-secreted factors in combinations of either a) TGF-α and TGF-β or b) BMP15 and GDF9 – resulting in the GLC-A and GLC-B clones, respectively. It has been shown that there exists a bi-directional relationship between the oocyte and follicular somatic cells, including GCs, and which regulate a diverse spectrum of granulosa cell function *in vitro* (Buccione et al., 1990; Vanderhyden et al., 1990). BMP15 and GDF9, as part of the TGF-β superfamily are necessary for early folliculogenesis and have been linked to ovarian dysfunction, and TGF-α has been identified as a major prospect for conferring the actions of oocytes in bovine GCs (Gilchrist et al., 2004; Glister et al., 2003; Juengel and McNatty, 2005; Jung et al., 2017; Shimasaki et al., 2004). In the present study, our differentiation protocol could successfully transition iPSCs into GLCs as reflected by the lower expression levels observed in RT-qPCR analysis of all pluripotency markers (NANOG, OCT4, SOX2, DPPA2, and DPPA4) in the GLCs. Furthermore, the GLCs displayed the typical spindle-shaped morphology (fibroblast-like) of human GCs and possessed high proliferative potential – both in subculture capacity and the ability for long-term cultivation of over thirty days. The post-differentiation stability and excellent growth characteristics of the GLCs suggest it could be a robust cellular source for future *in vitro* studies.

Anti-Müllerian hormone is also a member of the TGF-β superfamily, and in females AMH is secreted after birth by human GCs in the pre-antral and small antral follicles during early stages of folliculogenesis and is a prominent marker of granulosa cells (Depmann et al., 2015). Through paracrine signalling, AMH preserves the inactive state of a proportion of primary follicles by inhibiting the effect of follicle-stimulating hormone (FSH) on developing follicles (Nilsson et al., 2007; Sacchi et al., 2016). In XX mice AMH gene deletion led to early reduction of the ovarian reserve (Durlinger et al., 1999). Recently, missense mutations of AMH found in congenital hypogonadotropic hypogonadism (CHH) patients were associated with loss-of-function *in vitro* (Malone et al., 2019). The gonadotropin FSH is vital for follicular growth and Estradiol production in women, and the action of FSH is mediated by its receptor (FSH-R) that is expressed by GCs in the ovary (Desai et al., 2013). Deficiency of FSH-FSH-R interaction due to inactivating gene variants can cause polycystic ovary syndrome (PCOS) and infertility in women (Haqiqi et al., 2019; Laven, 2019). The elevated presence of both these important GC biomarkers in the GLCs relative to the control KGN cell line was confirmed by immunofluorescence of FSH-R and AMH ELISA measurement – which indicates an additional likeness of the GLCs to human granulosa cells.

To further assess the applicability of our GLCs as model for GCs, an extensive characterisation of the GLCs, KGN, and UP transcriptome was carried out based on scRNA-seq and RT-qPCR analysis of classical markers involved in GC fate (cell differentiation and proliferation) and function (folliculogenesis and steroidogenesis). Intriguingly, we observed UPs also expressed some of the genes analysed, which may be attributed to the known spontaneous differentiation potential of urine-derived cells such as UPs (Bharadwaj et al., 2013; Lazzeri et al., 2015; Liu et al., 2018; Wan et al., 2018). This phenomenon was also reflected in culture of our UPs during which a portion of cells within the UP population spontaneously differentiated into a mix of renal cell types. In granulosa cells, the gonadotropin-dependent enzyme cytochrome P450 aromatase (*CYP19A1*) catalyses the conversion of androgens into estrogens (Dunlop and Anderson, 2014) and was moderately expressed in the GLCs. As the most prevalent and potent form of endogenous Estrogen, estradiol (E2) is the primary steroid hormone synthesised by GCs within the ovarian follicles of premenopausal women (Ryan, 1982). In female sexual organs, E2 acts globally to regulate differentiation and function of reproductive tissues, drives cell cycle progression of GCs, and modulates folliculogenesis until ovulation (Baerwald et al., 2012; Chauvin et al., 2022; Quirk et al., 2006). Herein, the major functional determinant of the steroidogenic identity of the GLCs rests upon the production of E2, which was quantified by LC-MS/MS. Importantly, E2 was secreted in the growth medium of both GLC clones, although at slightly lower concentrations compared to KGN cells. This might be explained by the higher AMH levels in the GLCs relative to KGN – since it has been shown that a high amount of AMH in granulosa cells is concomitant with partial inhibition of FSH-induced E2 synthesis and leads to a decreased level of CYP19A1 expression (Chang et al., 2013).

Estradiol confers its effect mainly through its α and β receptor isoforms encoded by the Estrogen receptor 1 (*ESR1*) and *ESR2* genes, respectively (Taylor et al., 2010) – and mutations in these genes (particularly ESR2) have been linked in humans to ovulatory defects, primary ovarian insufficiency (POI), 46 XY VSDs, and granulosa cell tumours (Baetens et al., 2018; Ciucci et al., 2018; Lang-Muritano et al., 2018; Sundarrajan et al., 2001). The analysis of GC markers revealed ESR2 was indeed expressed in the GLCs whereas ESR1 was only slightly present, which is in congruency with the known differential expression of ESR1/2 in the ovary – with ESR2 mainly located in GCs throughout follicular development, while ESR1 is mostly found in theca cells and the surface epithelium (Coppens et al., 1993; Hillier et al., 1998; Scobie et al., 2002). In general, the tendency of granulosa marker expression in the GLCs was similar to the KGN control cell line, however the expression level of some markers – notably hydroxysteroid 17-beta dehydrogenase 1 (HSD17B1) was highly elevated in the GLCs. HSD17B1 is a steroidogenic enzyme crucial for the conversion of estrone to biologically active estradiol and is mainly expressed in the placenta and GCs of growing follicles in the ovary (Luu-The et al., 1995; Tremblay et al., 1989). It has been postulated HSD17B1 plays a key role in ovulation and is important in the pathophysiology and spread of estrogen-reliant breast carcinomas (Miller and Auchus, 2011; Sasano et al., 1996).

Another key regulatory factor for maintenance of the ovary in mammals is the highly conserved transcription factor *FOXL2* – one of the earliest expressed markers during ovarian differentiation (Cocquet et al., 2002; Veitia, 2010). Significantly, immunofluorescence characterization showed FOXL2 was clearly detectable in all GLC clones. The work of Uhlenhaut and co. showed the fundamental importance of FOXL2 in maintaining the ovarian phenotype throughout adulthood, by acting together with ESR1 to directly supress in an antagonistic manner a key male-promoting gene SRY-box9 (*SOX9*). Additionally, the study highlighted FOXL2 is essential for preserving granulosa cell identity in females as shown by FOXL2 knockout in GCs of mice resulting in testicular Sertoli cell marker expression and formations in the ovary resembling seminiferous tubules (Uhlenhaut et al., 2009). CYP19A1, that is involved in estrogen synthesis (as described before) is up-regulated by FOXL2, and FOXL2 regulates expression of Follistatin (*FST*) which plays an important role in ovarian follicle assembly (Caburet et al., 2012; Pannetier et al., 2006). Interestingly, FST was also highly expressed in the GLCs generated in this study. Functional testing *in vitro* has linked the loss of FOXL2 transcriptional activity in humans to Blepharophimosis, ptosis, and epicanthus inversus syndrome (BPES) of which Type I BPES is a form of POI (De Baere et al., 2001; Dipietromaria et al., 2009; Todeschini et al., 2011). Somatic point mutations of FOXL2 have also been associated with granulosa cell tumours in the ovary (Rosario et al., 2014; Shah et al., 2009; Weis-Banke et al., 2020) – and it is important to note the KGN cell line carries a form of FOXL2 with a variant linked to the loss of apoptotic function in GC tumours (Cheng et al., 2013).

Granulosa cell population type varies according to the stages of folliculogenesis (Gougeon, 2010): Initially primordial follicles are mobilised to become primary follicles, after which growing GCs (early-stage GCs) form multiple layers around the oocyte that in concert with a layer of theca cells initiate estrogen synthesis and develops into secondary/preantral follicles. At this time under stimulation of FSH antral follicles form (which contain mural GCs and cumulus cells). Then through selection between developing follicles only a few follicles progress to the preovulatory phase (that have late GCs) as the rest become atretic, followed by ovulation and finally luteinization of theca cells and GCs in the corpus luteum (Georges et al., 2014). In the scRNA-seq and RT-qPCR analysis we observed FOXL2 was expressed in the GLCs, but expression levels were moderately lower compared to KGN cells. Interestingly, FOXL2 is predominantly localized *in vivo* to undifferentiated ovarian GCs, with expression highest in GCs of primordial follicles which progressively decreases until formation of pre-antral/antral follicles and is not present at all in the corpus luteum (Hu et al., 2011; Kerr et al., 2013; Park et al., 2010; Park et al., 2014; Pisarska et al., 2004; Uda et al., 2004). With this in mind, the gene expression analysis of the GLCs revealed an expression profile primarily corresponding to mural GCs, yet also several marker genes of early-stage GCs and some markers of late GCs. This indicates dynamic cellular states within the overall cell population of the GLCs, which highlights the degree of differentiation between individual granulosa-like cells.

This could also explain the differences in expression of FOXL2 in the GLCs based on the granulosa cell-state of the specific follicular phase. This inherent differentiation ability and stemness that is characteristic of GCs has also been confirmed in previous studies investigating human gonadal niche cells – including GCs (Jozkowiak et al., 2020; Zhang et al., 2018). Hence, within this context we regarded the GLCs as a differentiating Granulosa cell model, which may potentially be an excellent cellular source to also study several developmental processes/phases of human folliculogenesis.

## CONCLUSION

Taken together, in this study we utilised for the first time UPs as alternative cell source to reprogram as iPSCs for rapid and efficient differentiation into GLCs, without compromising the original genome of source cells. The GLCs displayed characteristic morphology of human GCs and showed high proliferative potential during long-term culture. The GLCs recapitulated expression of classical granulosa-specific markers involved in human GC cell proliferation and differentiation, steroidogenesis, and granulosa dependent folliculogenesis. Considering that the GLCs are donor-derived it could serve as proxy for the protocol established in this study to be used for generation of patient-specific personalised GC-models to investigate mechanism of disease in VSDs. Moreover, the dynamic cellular state of the GLCs indicates a strong potential applicability as gonadal cell-model for ovarian development in women.

## ACKNOWLEDGEMENTS

This work was supported by the Schweizerischer Nationalfonds zur Förderung der Wissenschaftlichen Forschung, Grant Number: 320030_130645. We thank Valentin Braun and Christoph Seger (Dr. Risch Laboratories, Buchs SG, Switzerland) for their help in performing the Estradiol measurements.

## AUTHOR CONTRIBUTIONS

DH and AB-L conceived the project. DH performed the experiments, analysed the data and wrote the article. DRG assisted with data analysis and with some illustrations. AB-L supervised the research and provided final edits to improve the accuracy of the presentation.

