## Supplemental Table 1 for "Generation of a human ovarian granulosa cell model from induced pluripotent stem cells"

**Table S1. Primer sequences**

|  | **RT-qPCR** |
| --- | --- |
| **Target** | **Sequence 5`-3`** |
| ***PPIA*** | Fw: GGC AAA TGC TGC ACC CAA CAC A Rv: TGC TGG TCT TGC CAT TCC TGG A |
| ***NANOG*** | Fw: CTC CAA CAT CCT GAA CCT CAG C Rv: CGT CAC ACC ATT GCT ATT CTT CG |
| ***POU5F1 (OCT4)*** | Fw: CCT GAA GCA GAA GAG GAT CAC C  Rv: AAA GCG GCA GAT GGT CGT TTG G |
| ***SOX2*** | Fw: GCT ACA GCA TGA TGC AGG ACC A  Rv: TCT GCG AGC TGG TCA TGG AGT T |
| ***DPPA2*** | Fw: GGA CTG GTG TCA ACA ACT CGG T Rv: TCA CTG CCT TGC GTT TCC TCG A |
| ***DPPA4*** | Fw: CTC CAC AGA GAA GTC GAG GGA A Rv: GGT TGT CAG TGT GCT CTG CCT T |
| ***FOXL2*** | Fw: CGG AGA AGA GGC TCA CGC TGT Rv: CTG AGG TTG TGG CGG ATG CTA T |
| ***STAR*** | Fw: TAC GTG GCT ACT CAG CAT CGA C Rv: TCA ACA CCT GGC TTC AGA GGC A |
| ***INHBA*** | Fw: GGA TGA CAT TGG AAG GAG GGC A Rv: ACT GAC AGG TCA CTG CCT TCC T |
| **AMH** | Fw: CGC TGC TTC ACA CGG ATG ACC Rv: GGT GGC GAC TCC TCG AGT TCC |
| ***ZEB2*** | Fw: AAT GCA CAG AGT CTG GCA AGG C  Rv: CTG CTG ATG TGC GAA CTG TAG G |
| ***CD44*** | Fw: CCA GAA GGA ACA GTC GTT TGG C  Rv: ACT GTC CTC TGG GCT TGG TGT T |
| ***CYP19A1*** | Fw: GAC GCA GGA TTT CCA CAG AAG AG  Rv: ATG GTG TCA GGA GCT GCG ATC A |
| ***CYP11A1*** | Fw: TGG CAT CCT CTA CAG ACT CCT G Rv: CTT CAG GTT GCG TGC CAT CTC A |
| ***HSD17B1*** | Fw: TTC CTG CCA GAC ATG AAG AGG C Rv: AGA ACC GCC AGA CTC TCG CAT A |
| ***HSD3B2*** | Fw: CGC CTG TAT CAT TGA TGT CTT TGG Rv: CTG GTG TAG ATG AAG ACT GGC AC |
| ***NR5A1/ SF1*** | Fw: CCA GAC CTT CAT CTC CAT CGT G Rv: TGG CGG TAG ATG TGG TCG AAC A |
| **AR** | Fw: ATG GTG AGC AGA GTG CCC TAT C Rv: ATG GTC CCT GGC AGT CTC CAA A |
| ***ESR1*** | Fw: GCT TAC TGA CCA ACC TGG CAG A Rv: GGA TCT CTA GCC AGG CAC ATT C |
| ***ESR2*** | Fw: ATG GAG TCT GGT CGT GTC AAG G Rv: TAA CAC TTC CGA AGT CGG CAG |
| **PGR** | Fw: GTC GCC TTA GAA AGT GCT GTC AG Rv: GCT TGG CTT TCA TTT GGA ACG CC |
| **FSH-R** | Fw: GGT TTG TCC TCA CCA AGC TTC G Rv: GGT TGG AGA ACA CAT CTG CCT C |
| **LH-R** | Fw: GGA GAA GAT GCA CAA TGG AGC C Rv: CGT GGC AAT TAG CCT CTG AAT GG |
