## Supplemental Figure 1 for "Generation of a human ovarian granulosa cell model from induced pluripotent stem cells"

### Estradiol quantification

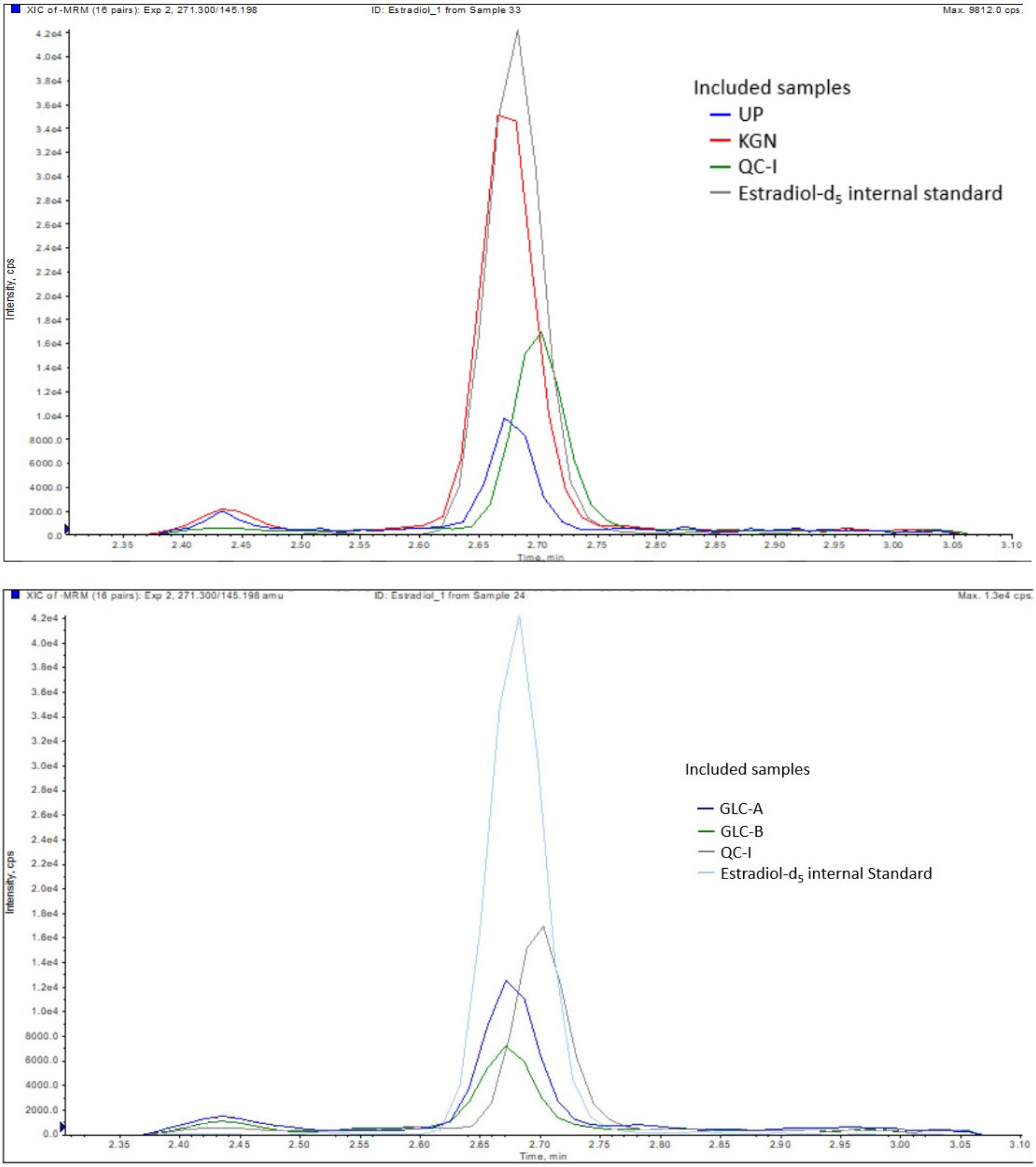


**Figure S1. Representative chromatograms of the LC-MS/MS analysis of Estradiol.** Overlays are of Estradiol quantifier ion traces for UP and KGN cell samples (top) and cell samples of the GLCs (bottom) with the internal standard E2-d$_5$. Note: Quantifier ion traces are represented as raw readings and not yet normalised for total cell count in samples. CPS, counts per second; QC-I, quality control-internal.
